# Role of Tyrosine Phosphorylation in PTP-PEST

**DOI:** 10.1101/2023.11.28.569137

**Authors:** T S Sreevidya, Amrutha Manikandan, N Manoj, Madhulika Dixit, Satyavani Vemparala

**Affiliations:** The Institute of Mathematical Sciences, C.I.T. Campus, Taramani, Chennai 600113, India; Homi Bhabha National Institute, Training School Complex, Anushakti Nagar, Mumbai 400094, India; Department of Biotechnology, Indian Institute of Technology Madras, Chennai 600036, India

## Abstract

We study the influence of tyrosine phosphorylation on PTP-PEST, a cytosolic protein tyrosine phosphatase. Utilizing a combination of experimental data and computational modeling, specific tyrosine sites, notably Y64 and Y88, are identified for potential phosphorylation. Phosphorylation at these sites affects loop dynamics near the catalytic site, altering interactions among key residues and modifying the binding pocket’s size. This, in turn, impacts substrate binding, as indicated by changes in binding energy. Our findings provide insights into the structural and functional consequences of tyrosine phosphorylation on PTP-PEST, enhancing our understanding of its effects on substrate binding and catalytic conformation.

## I. INTRODUCTION

The regulation of protein activity is crucial for controlling cell behavior and maintaining normal cellular function. This regulation occurs at different levels, including transcriptional, translational, and post-translational modifications (PTMs)^1,2^. Among PTMs, protein phosphorylation is a rapidly reversible modification that plays a significant regulatory role in various aspects of cell physiology^3–6^. Phosphorylation and dephosphorylation, catalyzed by protein kinases and protein phosphatases, can modify protein function in multiple ways. They can increase or decrease biological activity, stabilize or mark proteins for degradation, facilitate or inhibit movement between different parts of the cell, and initiate or disrupt protein-protein interactions ^5–12^. In eukaryotes, phosphorylation primarily occurs on serine, threonine, and tyrosine residues^13,14^. Although tyrosine phosphorylation represents a small fraction of the phospho-proteome (less than 2%)^15–17^, it plays a crucial role in regulating various processes in eukaryotic cells, including cell signaling, growth, proliferation, differentiation, migration, survival, and apoptosis ^11,16,18–27^.

PTP-PEST is an 88 kDa cytosolic protein and a member of the protein tyrosine phosphatase (PTP) family, which represents the largest group of phosphatase genes^25,28,29^. It comprises of an N-terminal catalytic domain that features the active site signature motif C(X)5R^3^ and is characterized by the presence of multiple Proline-Glutamic acid-Serine-Threonine (P-E-S-T) motifs ^29–32^. By dephosphorylating multiple substrates^33^, which include HER2, FAK, PYK2, PSTPIP, WASP, p130Cas, and paxillin, PTP-PEST assumes a pivotal role in the regulation of diverse physiological processes. These encompass cell adhesion, cell migration, immune response, embryonic development, and apoptosis^25,28,34,35^.

PTPs share a common two-step catalytic mechanism, facilitated by a well-organized active site with specific residues. In PTP-PEST, the active site consists of five major loop regions: the phosphate-binding loop (P-loop), WPD-loop, Q-loop, pTyr-loop, and E-loop, numbered according to the PTP-PEST sequence (See Figure 1)^36^. The P-loop, which contains the PTP active site motif C(X)5R, plays a crucial role in catalysis. It includes the catalytic cysteine (C231) and the invariant arginine (R237). The guanidinium group of R237 coordinates the phosphate group during substrate binding and the formation of a phospho-cysteine intermediate. The WPD-loop harbors the catalytic acid/base aspartate (D199), which acts as a general acid to donate a proton to the leaving group and later as a general base to remove a proton from a water molecule. This process facilitates hydrolysis of the phosphorous-sulfur bond, resulting in the release of free phosphate. The WPD-loop can adopt both closed (active) and open (inactive) conformations and serves as a flexible gate to the active site^37,38^. The Q-loop contains conserved glutamine residues (Q278, Q282) that coordinate a water molecule essential for hydrolysis. The pTyr-recognition loop (pTyr-loop) contains the conserved motif “KNRY” (K61-Y64), which provides specificity for phosphorylated tyrosine (pTyr). The conserved tyrosine residue (Y64) forms aromatic *π − π* interactions with the substrate’s pTyr residue, facilitating substrate binding to the active site^36,38^. The E-loop (putatively spanning R134 to K142) contains multiple conserved residues that appear to coordinate the dynamics of the WPD-loop. The conserved glutamate (E137) in the E-loop forms a bi-partite hydrogen bonding interaction with the side chain of the P-loop’s arginine (R237), while the conserved lysine (K142) of the E-loop often forms a hydrogen bond with the catalytic aspartate (D199) of the WPD-loop in its closed conformation. All the residues mentioned thus far are critical for the essential functioning of PTP-PEST^36^.

**FIG. 1.**
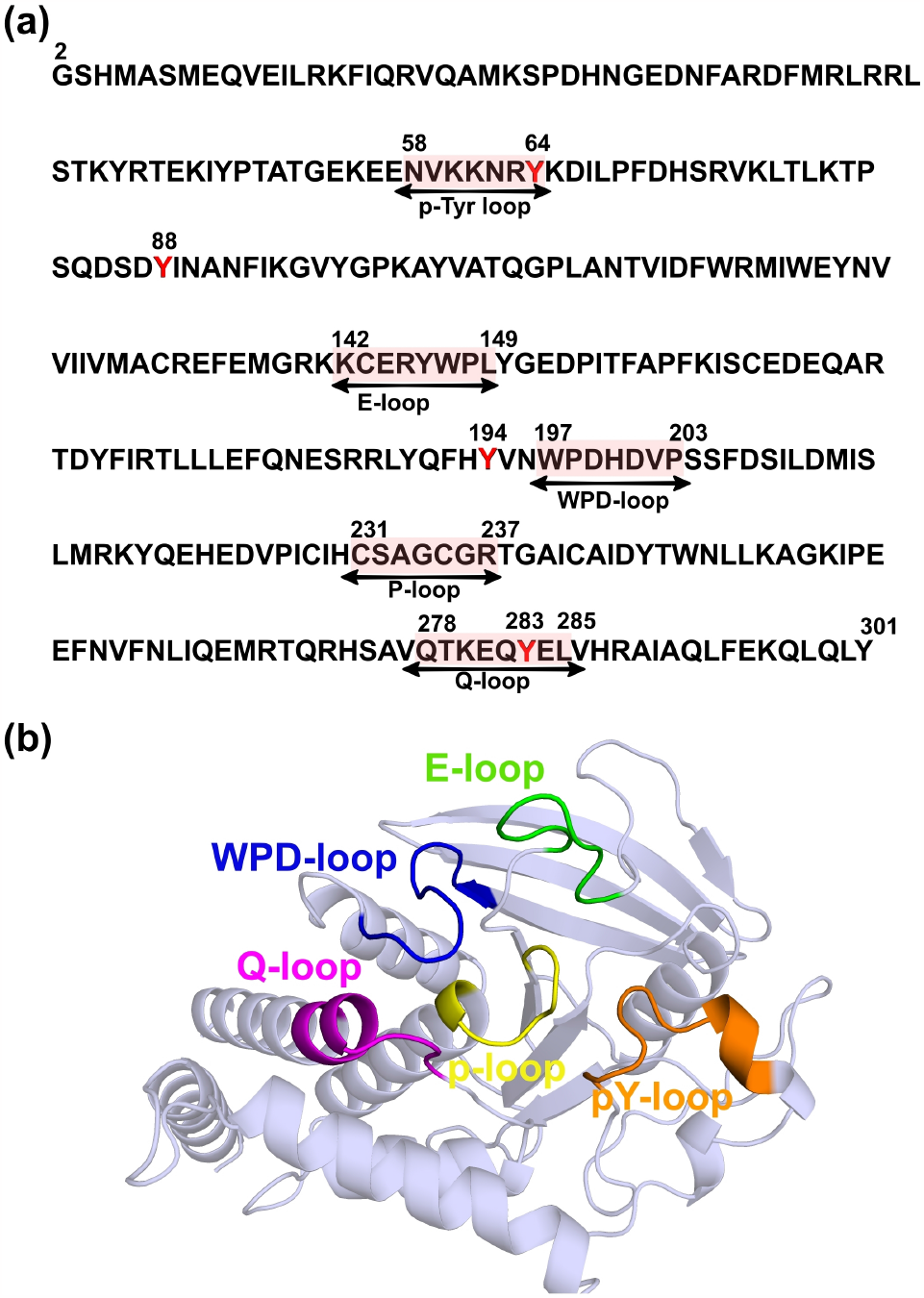
(a) Sequence of PTP-PEST protein (PDB ID: 5HDE). Loops forming the catalytic site are highlighted in red along-with numbering. Phosphorylated tyrosines are shown as red text. (b) Cartoon representation of PTP-PEST(PDB ID : 5HDE, loops involved in catalysis are highlighted. P-loop, pY-loop, WPD-loop, Q-loop and E-loop are shown as yellow, orange, blue, magenta and green cartoon representations respectively.

The potential for reversible phosphorylation to regulate the activity of PTP-PEST has been investigated in earlier studies^29^. The serine phosphorylation of PTP-PEST at S39 and S19 has been observed to modulate substrate affinity and enzymatic activity^33,39^. However, the N-terminal phosphatase domain of PTP-PEST also has a large number of tyrosine residues and the present study is aimed at understanding (1) is tyrosine phosphorylation possible in PTP-PEST and can it be shown experimentally? (2) can specific tyrosine phosphosites be identified by bioinfomatic tools? (3) what are the implications of tyrosine phosphorylation at the predicted phosphosites on the structure and catalytic binding pocket conformation via detailed MD simulations? (4) how does binding of substrates gets affected by such tyrosine phosphorylation via docking studies?

## II. METHODS

### A. Protein Purification and Cell Culture

The catalytic domain of PTP-PEST (1-300 amino acids) was cloned into the pET28a(+) bacterial expression vector and purified as described previously^40,41^. Briefly, plasmids were transformed into BL21(DE3)-RIL competent cells and protein expression was induced in a 2.4L culture grown at 37°C, 180 rpm at an O.D of 1 using 150*μ*M IPTG. The culture was further grown for 16 hours, 180rpm at 16°C. Bacterial pellets were lysed in a buffer containing 50 mM Tris, 200 mM NaCl, 1 mM PMSF, 0.5 mM DTT, and 1 mM imidazole^40,41^. The bacterial suspension was sonicated for 5 minutes at 35% amplitude (pulse at 5s ON and 5s OFF). The cell suspension was then pelleted at 8000 rpm for 90 minutes at 4°C after which the supernatant was loaded onto a Ni^2+^ -NTA resin containing column and allowed to flow through the resin under gravity flow at 4°C. The column was washed, and the pure protein was eluted using a buffer containing 200 mM imidazole. After purification, the protein was desalted using a buffer exchange method with a 10 kDa molecular weight cut-off Millipore concentrator. Finally, the protein was resuspended for storage according to the previously described protocol^40,41^. EA.hy926 cells (CRL-2922) were cultured in DMEM high glucose media (Himedia, catalog number: AT007) supplemented with sodium bicarbonate and 10% Fetal Bovine Serum (South Americal origin, MP Biomedicals). Cells were grown till a confluent monolayer, following which they were used for experimentation.

### B. In-vitro kinase assay and Immunoprecipitation

EA.hy926 cells were washed twice with 1X PBS. A non-denaturing lysis buffer containing tyrosine phosphatase inhibitors (50mM Tris, 150mM NaCl, 1% Triton X100, 1X protease inhibitor cocktail and 2mM Sodium orthovanadate) was added to the dish and incubated on ice. The cells were scraped after incubation on ice and the harvested cells were transferred into freshly autoclaved tubes. The tubes were vortexed at maximum speed for 30s and transferred back on the ice for one minute. After repeating this step 5 times, the cells were sonicated for 2 minutes with a pulse of 2s ON and 8s OFF, at 30% amplitude. The sonicated cells were centrifuged at 12000 rpm for 10 minutes at 4°C. An in-vitro kinase assay was performed to assess the phosphorylation of bacterially purified WT PTP-PEST by kinases from the cell lysate. Protein estimation was performed using Bradfords reagent after which 100*μ*g of purified WT PTP-PEST protein was incubated with 100*μ*g of cell lysate and 500*μ*l of 1X Kinase buffer (125mM HEPES pH 7.4, 30mM MgCl_2_, 2mM MnCl_2_, 0.2mM DTT). The reaction was supplemented with 200 *μ*M of ATP from a stock of 10mM ATP. The reaction was incubated at 30°C in a water bath for 30 minutes after which the tubes were placed on ice and EDTA was added to a final concentration of 5mM to stop the reaction. Pull down of PTP-PEST protein was performed using Ni^2+^-NTA beads, after which tubes were incubated overnight on a rotospin at 4°C at 12 rpm. Samples were washed three times with wash buffer containing 50mM Tris-Cl, 150mM NaCl and 20mM imidazole. Proteins were eluted from the beads by the addition of 30*μ*l of 2X loading dye containing *β*-mercaptoethanol and heating at 95°C for 5 minutes. The elute was loaded on a 10% SDS PAGE gel for Western Blotting. Membranes were immunoblotted for total-phosphotyrosine (Cell Signaling Technology No. 8954) and PTP-PEST (Cell Signaling Technology No. 8954) respectively as per manufacturers instructions.

### C. Bioinformatics analysis of tyrosine phosphorylation sites on PTP-PEST

Prediction algorithms and database mining methods were both used to determine putative tyrosine phosphorylation sites on PTP-PEST. The NetPhos 3.1 webserver which uses neural networks to identify phosphosites by providing a phosphorylation potential score^42^, was used for the prediction of tyrosine phosphorylation sites. The primary amino acid sequence of PTP-PEST was used as input for NetPhos 3.1 predictions and only those tyrosines that scored above a high threshold of 0.7 were chosen for further analysis. The GPS 6.0 webserver^43,44^ was also used to predict putative phosphorylation sites. This tool predicts phosphorylation sites based on three integrated machine learning approaches and also identifies putative kinases that may phosphorylate a given residue. The highest threshold was used to filter GPS 6.0 predictions after providing the primary amino acid sequence of PTP-PEST as the input. Database mining was performed using the Eukaryotic Phosphorylation Site Database (EPSD)^45^. This database curates open-source high throughput phosphoproteomics data from different studies and integrates datasets from 13 different phosphorylation site databases. Consurf analysis was performed to determine the evolutionary significance of the putative phosphorylation sites^46^. The Consurf webserver uses different sequence alignment methods to estimate the evolutionary conservation of a particular amino acid and assigns a score to it^45^. In case of the catalytic domain of PTP-PEST, the PDB ID 5HDE was used as the input, as ConSurf uses the crystal structure information to map evolutionarily conserved residues directly on the structure of the protein^47^. For the C-terminal domain of PTP-PEST, the primary amino acid sequence of this region was used as the input.

### D. Molecular Dynamics Simulation

The starting structure for control simulations of PTP-PEST is taken from Protein Data Bank: PDB ID 5HDE. Other systems with relevant tyrosines phosphorylated were created from this initial structure. While the catalytic cysteine (C231) in PTP-PEST is in a phosphorylated state in the crystal structure, the same was unphosphorylated in all the simulations here to mimic the wild-type. The correct protonation states of the ionizable residues were determined at the experimental pH of 7.2 using DEPTH server^48^ and appropriately changed before solvating the system. Each system was solvated using TIP3P water model^49^ and for overall charge neutralization and to mimic the physiological salt concentration, 0.15 mol/L NaCl salt was added. All the simulations were performed using Charmm36 forcefield^50,51^ in NAMD 2.12 software^52^. Energy minimization was done for 5000 steps using conjugate gradient method^53^. Langevin thermostat^54^ was used to maintain constant temperature at 300K. Nose-Hoover Langevin piston^55,56^ with a decay period of 100fs and a damping time of 50fs was used to maintain pressure at 1atm. A cut-off distance of 12°*A* was used to calculate short-range van der Waals (VDW) interactions and the long-range electrostatics interactions was calculated by the Particle Mesh Ewald(PME) method^57,58^. VMD software^59^ and the PyMOL Molecular Graphics System^60^ were used for visualization and analysis was done using in-house analysis codes using TCL scripting language and VMD plugins.

### E. Community Network Analysis

Community network analysis, conducted through the NetworkView plugin in VMD^61^, identifies connections between amino acids in proteins based on molecular dynamics trajectories. The analysis defines a network with nodes representing C_*α*_ atoms, connected by edges. Edges are formed if corresponding C_*α*_ atoms are within 4.5°*A* proximity for at least 75% of frames, excluding those with adjacent residue numbers. Edge weights are determined using the correlation matrix (*C*_*ij*_) data, employing the formula *w*_*ij*_ = *−* log(|*C*_*ij*_|). Calculated by Carma software^62^, correlation matrices indicate residue motion correlation, serving as a measure of information transfer. Community detection analysis, executed with gncommunities software, was applied to all systems in VMD, as detailed in the references^61,63,64^.

### F. Principle Component Analysis

To tackle the challenges of analyzing large datasets, dimensionality reduction techniques like Principal Component Analysis (PCA) are often employed and reveals insights into conformational differences^65–67^. PCA involves computing a covariance matrix from the dataset, followed by eigenvalue decomposition. The resulting eigenvectors, known as principal components, capture the most significant information, with eigenvalues indicating variance explained. In our study, PCA was applied to an M x N matrix, representing frames and C*α* atoms, respectively. PC1, derived from this analysis using the bio3d package in R^68^, represents the axis of maximum variance in structure distribution.

### G. Docking Analysis and Binding Energy Calculations

Docking studies were conducted using AutoDock Vina 1.1.2 software^69^ for the HER2-derived peptide and SRC-derived peptide with all the PTP-PEST systems. Residues known from literature to be important for substrate recognition and binding (Y64, E137, R140, R141, K142, W197, P198, D199, C231-R237, Q275, Q282) were selected. The center of mass of these residues was measured, and the coordinates were used to define the center of the grid for active-site docking. A cubic box with dimensions of 30 × 30 x 30 points was created around this center for the docking simulations. Ten bound ligand conformations were obtained from the docking simulations and visualized in PyMOL^70^ along with the protein. Conformations where the phosphate group was facing inward towards the binding groove were selected for further analysis. The binding affinities of the selected conformations with their respective protein partners were calculated using the PRODIGY web server^71,72^. PRODIGY calculates the binding affinity based solely on the structural properties of the protein-protein complexes. It calculates the number of intermolecular contacts (ICs) at the interface within a threshold distance of 5.5°*A* and lists them according to their contact properties. The binding affinity is determined by combining this information with properties of the non-interacting surfaces (NIS).

## III. RESULTS

### A. The catalytic domain of PTP-PEST undergoes tyrosine phosphorylation

An in-vitro kinase assay was performed on bacterially purified PTP-PEST using mammalian cell lysate. Since tyrosine phosphorylation is transient and constitutes only 1% of the total phosphorylation events in the cell, the in-vitro kinase assay was conducted in the presence of 200*μ*M of ATP to increase the pool of phosphorylated PTP-PEST. As depicted in Figure 2, upon treatment with mammalian cell lysate and immunoprecipitation of the His-tagged catalytic domain using Ni^2+^-NTA beads, there was a clear spike in the total tyrosine phosphorylation of the bacterially purified protein, whereas no bands were observed in the absence of cell lysate (Figure 2).

**FIG. 2.**
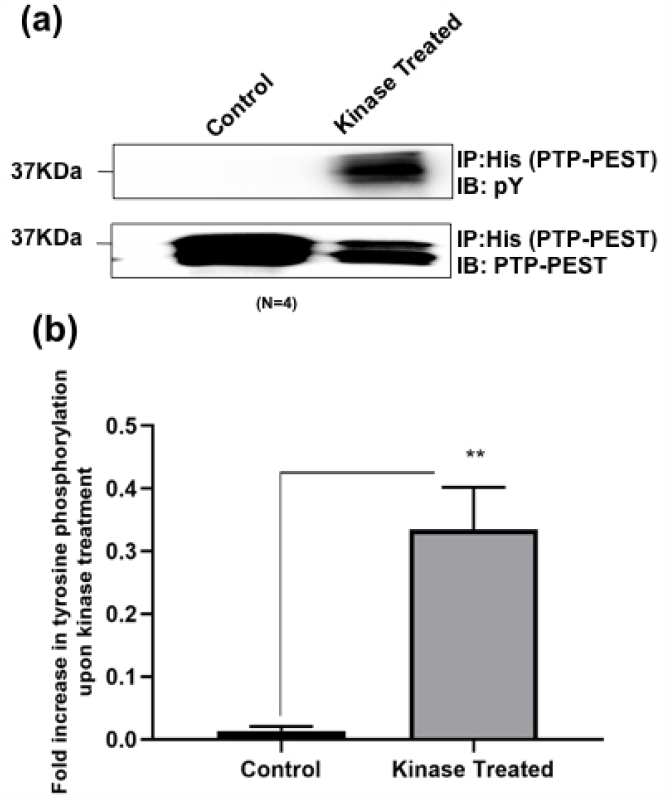
(a) Representative western blot showing immunoprecipitation of 100*μ*g of PTP-PEST after in-vitro kinase assay at 30°C upon incubation with kinase buffer and 100*μ*g of EA.hy926 cell lysate (b)The bar graph summarizes data from five independent experiments as mean increase in total tyrosine phosphorylation of PTP-PEST in the absence or presence of in-vitro kinase assay using endothelial cell lysate, normalized to the amount of PTP-PEST pulled down.

Following experimental confirmation that the catalytic domain of PTP-PEST can undergo tyrosine phosphorylation, we mapped putative phosphorylation sites on the crystal structure of the catalytic domain. Figure 3(a) shows all the tyrosine residues in the N-terminal phosphotase domain that were considered for further analysis via bioinformatic tools to identify the best possible tyrosine residues that are potential phosphosites. The location of these tyrosine residues with respect to functionally relevant loops is shown in Figure 3(b). Using bioinformatics tools NetPhos3.0^73^ and GPS6.0^43^, along with the Eukaryotic Phosphorylation site Database (EPSD)^45^, we found predicted and known PTP-PEST phosphotyrosines primarily in the catalytic domain. Analysis of residue conservation and structure (Figure 3 and Table I) revealed that Y64 and Y88, located in the pY and *β*1-*β*1 loops respectively, were highly conserved and identified by both algorithms and curated in the database (Table I). Interestingly, mutating Y64 to Y64A significantly reduces PTP-PEST catalytic activity^33^. Y88 resides on the structurally plastic *β*1-*β*1 loop of PTP-PEST, speculated to involve substrate recognition site formation^33^. We also selected Y194, due to proximity to the WPD loop, and Y283, having high ConSurf score, as potential sites. Given their structural significance and high phosphorylation potential, Y64, Y88, Y194, and Y283 were chosen for further MD simulation analysis.

**TABLE I.** Analysis of tyrosine phosphorylation sites across PTP-PEST : The ✓ symbol represents prediction by the particular algorithm/ presence in database whereas - denotes absence. Consurf Scores are derived from the multiple sequence alignment, where scores between -0.161 to -1.449 are assigned based on increase in residue conservation and scores from - 0.161 to 3.148 are assigned based on decreased conservation of that residue.

| Y-Position | NetPhos | GPS | Database | ConSurf Score |
| --- | --- | --- | --- | --- |
| 42 | - | ✓ | ✓ | 1.424 |
| 48 | - | ✓ | ✓ | 0.624 |
| 64 | ✓ | ✓ | ✓ | -0.956 |
| 88 | ✓ | ✓ | ✓ | -1.378 |
| 103 | - | ✓ | - | -0.812 |
| 146 | - | ✓ | ✓ | -1.438 |
| 150 | - | ✓ | ✓ | 2.632 |
| 194 | ✓ | - | - | -0.300 |
| 219 | - | ✓ | - | 1.161 |
| 283 | ✓ | ✓ | - | -1.243 |
| 301 | - | - | ✓ | -0.828 |
| 387 | ✓ | ✓ | - | -1.002 |
| 560 | - | - | ✓ | 1.258 |
| 733 | - | ✓ | ✓ | 1.346 |

**FIG. 3.**
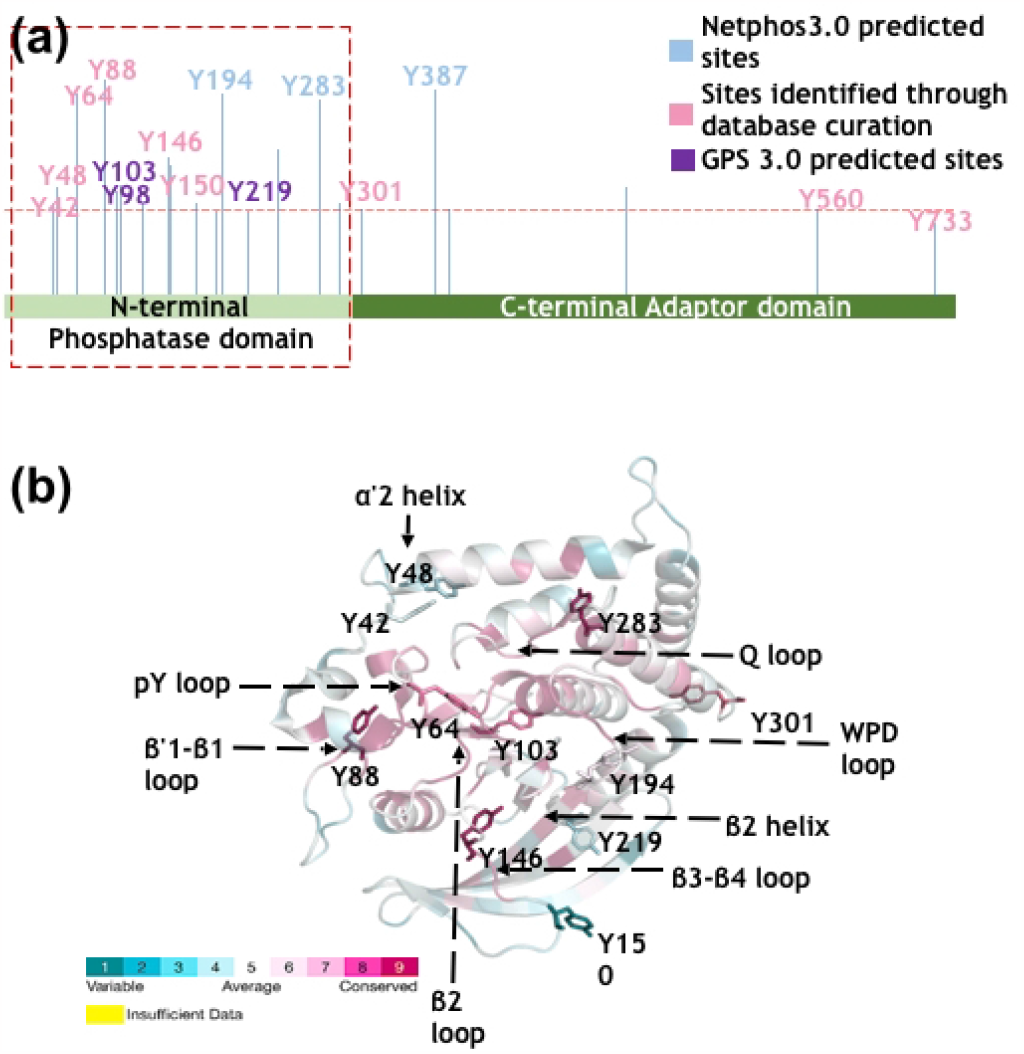
(a) Key tyrosine residues identified as possible phosphorylation sites via prediction algorithms and database mining methods. (b) Cartoon representation of 5 loops of PTP-PEST which are a part of the active site. P-loop, pY-loop, WPD-loop, Q-loop and E-loop are shown, coloured by the conserved residues. The tyrosines of interest are shown by stick representation.

### B. Global effects of phosphorylation of different tyrosine residues

Five systems were constructed in this study for detailed MD simulations to understand the effect of phosphorylation on key tyrosines identified earlier: wild type PTP-PEST (PDB ID: 5HDE), PTP-PEST phosphorylated at Y64, PTP-PEST phosphorylated at Y88, PTP-PEST phosphorylated at both Y64 and Y88, and PTP-PEST phosphorylated at Y64, Y88, Y194, and Y283. These systems are referred to as the control, pY64, pY88, pY64+pY88, and pY-all systems, respectively. While most analyses focus on the pY64 and pY88 systems, comparisons with the pY-all system are also performed and indicated accordingly. The details of the simulated systems are found in Table II.

**TABLE II.**
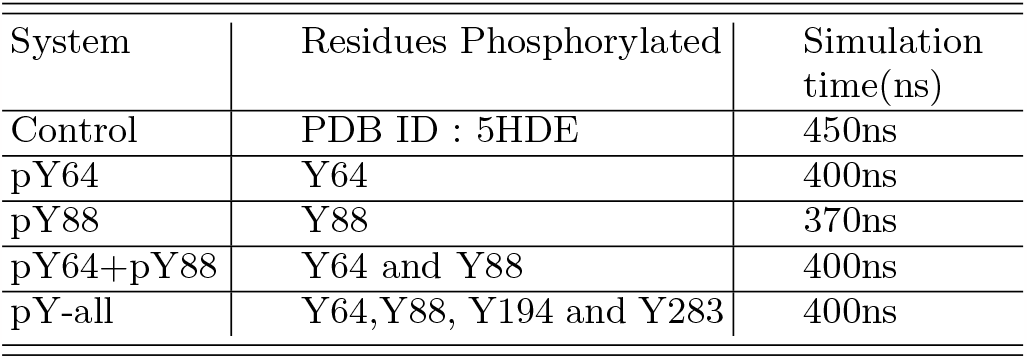
Simulation details of the systems studied in this study.

| System | Residues Phosphorylated | Simulation time(ns) |
| --- | --- | --- |
| Control | PDB ID : 5HDE | 450ns |
| pY64 | Y64 | 400ns |
| pY88 | Y88 | 370ns |
| pY64+pY88 | Y64 and Y88 | 400ns |
| pY-all | Y64, Y88, Y194 and Y283 | 400ns |

Figure 4 illustrates the positioning of the relevant tyrosine residues whose phosphorylation is pertinent to this study, along with important loops and residues involved in catalysis in PTP-PEST. The tyrosine residues that are putatively phosphorylated are all located on or proximal to the functional groups. Two tyrosines, Y64 and Y283, are situated on the periphery of PTP-PEST, while Y194 and Y88 are buried phosphosites. It can be envisioned that the charged residues on the functional loops may have their conformations significantly affected by the phosphorylation of tyrosines due to the introduction of net negative charges in their vicinity, leading to electrostatic interaction-induced conformational changes of the functional loops. The overall structural properties of PTP-PEST, with and without phosphorylation, were monitored via the time evolution of RMSD, RMSF, solvent-accessible surface area (SASA), and radius of gyration (*R*_*g*_). The phosphorylation of relevant tyrosines contributes to an increase in SASA values, as well as RMSD values (Figure S1(a,c)), suggesting increased local exposure to solvent via conformational changes on phosphorylation. The mobility of several residues is also affected by tyrosine phosphorylation (Figure S1(b)), further indicating local residue-level changes. The *R*_*g*_ data, however, does not show significant changes with the phosphorylation of tyrosine residue(s), suggesting that local phosphorylation of the tyrosine residues may not affect the global size of PTP-PEST (Figure S1(d)).

**FIG. 4.**
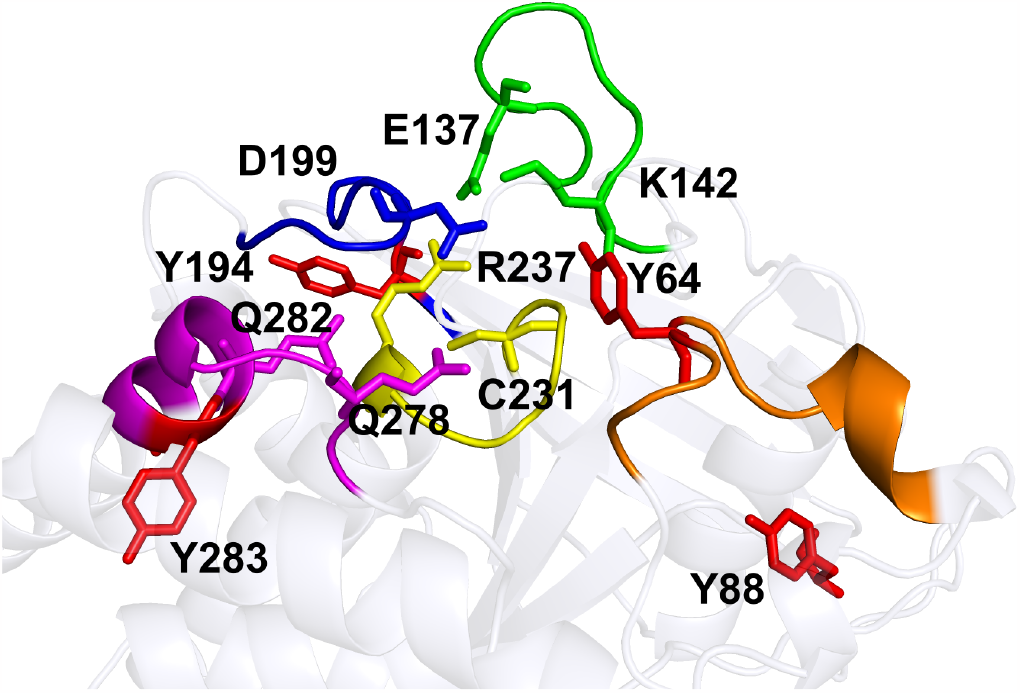
Cartoon representation of 5 loops of PTP-PEST which are a part of the active site. P-loop, pY-loop, WPD-loop, Q-loop and E-loop are shown in yellow, orange, blue, magenta and green cartoon representations respectively. The tyrosines of interest and other key residues of each loop are shown by stick representation.

To investigate the effect of phosphorylation on the residue-residue interactions within the protein, protein network analysis was performed. The protein network is perceived as a set of nodes (*C*_*α*_ atoms) connected by edges if the distance between the two residues is within 4.5 for at least 75% of the frames analyzed. The network consists of nodes and edges, forming substructures or communities of nodes that are more densely inter-connected with each other than with other nodes in the network. The number of communities (*N*_com_) indicates sub-regions of protein with relatively independent dynamics, but with multiple interaction pathways within the community. The goal is to investigate whether phosphorylation of the tyrosines alters the protein interaction network either via formation of new edges (new interactions) in the network, changes in interconnectivities between residues, or a change in the number of communities (*N*_com_). The results of such protein network analyses, seen in Figure 5, show that phosphorylations lead to a change in the *N*_com_ of all four phosphorylated systems (Figure 5). The number of distinct communities in the Control system is *N*_com_=11 (Figure S2). However, there is a decrease in the number of communities in the pY64 (*N*_com_=8), pY88 (*N*_com_=7), and pY64+pY88 (*N*_com_=7) systems (Figures 5(b-d)), while the number of communities increased in the pY-all (*N*_com_=13) system compared to the Control system. The decrease in number of communities indicates increased intra-protein connectivity, coalescing the overall number of highly-interacting protein residues. From the network analyses, phosphorylation of all relevant tyrosines (pY-all system) retains the original connectivity of the unphosphorylated system, suggesting cancellation effects.

**FIG. 5.**
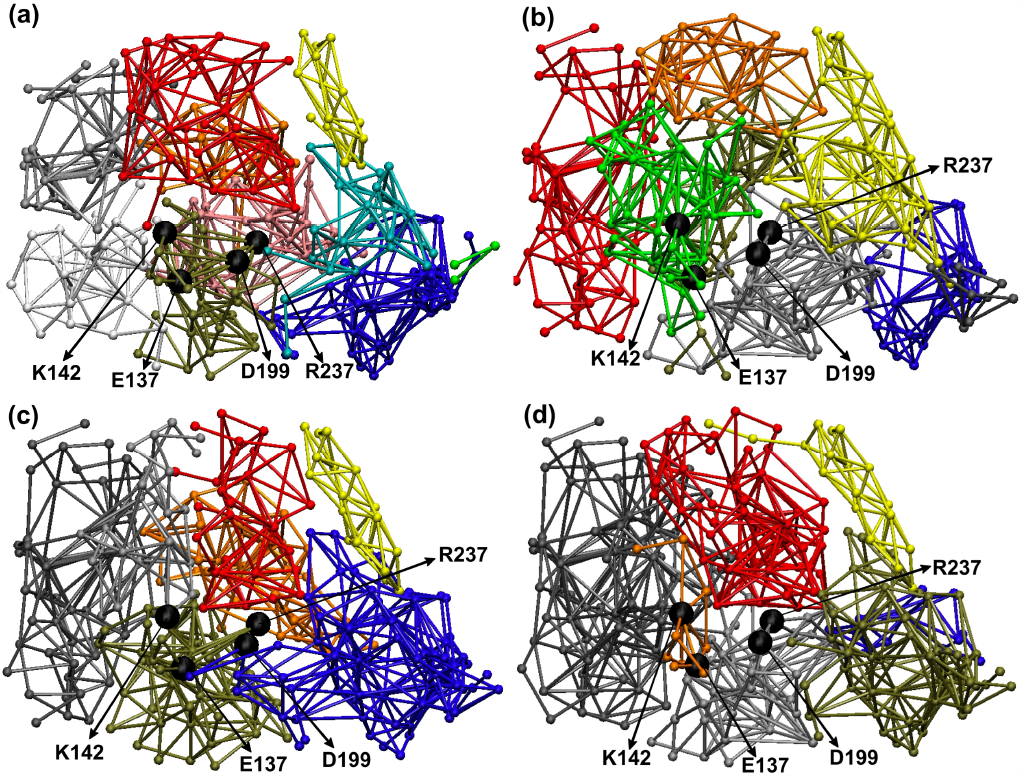
Network analysis over the last 50ns of (a) control (b) pY64 (c) pY88 (d) pY64+pY88 PTP-PEST systems

We further examine the effect of phosphorylation on two crucial catalytic loops, the P-loop and WPD-loop, and the functionally relevant residues in these loops (W197, D199, K142, E137, R237) regarding their presence in the network communities discussed above. We particularly focus on the P-loop (C231 to R237) and WPD loop (W197 to D199) as together they constitute the active site and the lid or gate to the active site. In the Control system, the P-loop is distributed across four communities: olive green (residues 231, 232, 237), red (residues 233, 235), orange (residue 234), and pink (residue 236). Post-phosphorylation, a reorganization of these residues was observed, leading to the P-loop being associated with fewer communities in the phosphorylated systems compared to Control. In pY64, the P-loop is in three communities: green (residues 231-234), yellow (residues 235, 236), and grey (residue 237). In pY88, it is in red (residues 231, 233, 234, 235, 236), grey (residue 232), and olive green (residue 237). In pY64+pY88, it is associated with red (residues 231, 233, 234, 235, 236), orange (residue 232), and grey (residue 237). Finally, in pY-all it is split between dark grey (residues 231, 233, 234, 235, 236, 237) and grey (residue 232). Unlike the P-loop, the WPD-loop residues (W197-D199) reside in the same community in all five systems, differing in which community and interacting residues. In Control, WPD loop is with K142, E137, R237 (olive green), while in pY64, K142 and E137 move to the green community with R237. In pY88 and pY-all, the WPD-loop is in a different community compared to K142, E137 and R237. In pY64+pY88, the WPD loop is with R237 (grey). The results suggest changes in strongly interacting communities and interactions between catalytically relevant loops and functionally relevant residues upon phosphorylation. The network analysis results also correlate with the cross-correlation analyses in Figure S2, shedding light on local correlations and distant protein interactions. As highlighted, tyrosine phosphorylation significantly mitigates intra-residue correlations, agreeing with network picture changes. Particularly, increased local correlations when all tyrosines are phosphorylated (Figure S3(e)) agree with the increased interaction communities. Together, these observations suggest phosphorylation(s) significantly affect key PTP-PEST residues, important for substrate recognition and binding.

### C. Role of tyrosine phosphorylation on local interactions

In the previous section, network analyses revealed phosphorylation of relevant tyrosine residues leads to differences in the protein network communities containing functionally relevant residues. Here, we probe how this change manifests in local interactions within catalytically important residues. A functionally important interaction network exists between key PTP-PEST residues crucial for catalysis: D199, K142, E137, and R237. These residues belong to the catalytic loops of PTP-PEST (Figure 4). D199 (WPD loop) acts as the catalytic acid/base for substrate hydrolysis, K142 (E-loop) hydrogen bonds with D199, E137 (E-loop) forms a bipartite H-bond with R237 (P-loop), and R237 guanidinium group coordinates the phosphate group during substrate binding and phospho-cysteine intermediate formation^36^. In the Control network, D199 interacts with K142 and R237, R237 with D199 and E137, K142 with D199 and E137, and E137 with K142 and R237. This strong intra-interaction manifests in their shared community in Control system(Figure S2), while after phosphorylation(s) they no longer share a community (Figure ref-fig:network(a)). This indicates phosphorylation of any of the 4 Ys changes interactions between these residues, resulting in different communities.

To further understand the effect of pY64, pY88, pY64+pY88 and pY-all on the mini-interaction network of 4 important PTP-PEST residues, distance analysis was done. Y64 is included in all the analyses as it determines binding pocket depth. In pY64, E137-R237 and D199-R237 interactions were retained (Figure S4(a)) while E137-K142 and D199-K142 were lost (Figure 6 (c,d)). This may be because a new pY64-K142 interaction from phosphate addition causes K142 to move, disrupting its E137 and D199 interactions (Figure 6(a,d)). In pY88, E137-K142, D199-K142 and D199-237 interactions were lost (Figure 6(a-c)) and only E137-R237 was retained. This may be from D199 and K142 moving from initial conformations (Figure S4(c)), enabling a new K142-Y64 interaction in pY88 (Figure 6(d)). Interestingly, pY64+pY88 effects resemble pY64 (Figure S4(a,c), Figure 6). In pY-all, only E137-R237 interaction was fully retained. Similar to pY64, E137-K142 and D199-K142 interactions were lost in pY-all (Figure 6(a,b)), possibly from the new pY64-K142 interaction disrupting K142’s E137 and D199 interactions (Figure S4(d)). In pY-all, the D199-237 interaction was lost at 170ns, regained at 270ns, and otherwise retained (Figure 6(c)). Changes in this key residue interaction network may impact substrate binding in the phosphorylated PTP-PEST systems.

**FIG. 6.**
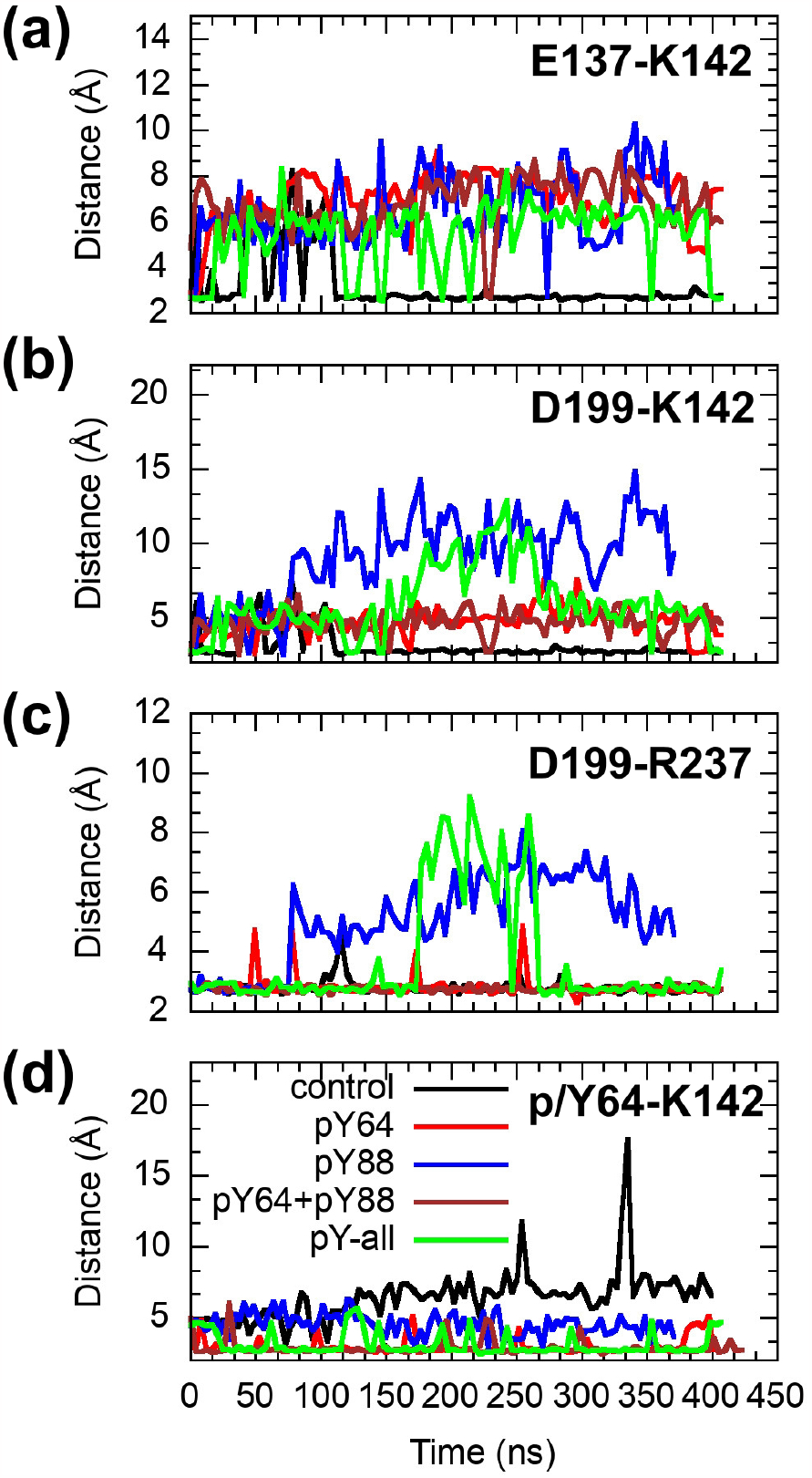
Time evolution of distances between (a) E137-K142 (b) D199-K142 (c) D199-R237 and (d) p/Y64-K142 over the length of the simulation.

### D. Catalytic pocket and lid dynamics affected by tyrosine phosphorylation

The WPD-loop acts as a flexible gate to the active site, observed in closed (active) and various open (inactive) conformations^3^. In classical PTPs, the tryptophan is an essential hinge residue important for loop flexibility^3^. Here, we explore the effect of tyrosine phosphorylation on both WPD dynamics and catalytic site access. Figure S5 shows the time evolution of Root Mean Square Deviation (RMSD) of residues 197-203 (WPD-loop) for all systems. The WPD loop undergoes conformational change in Control over the simulation timescale. The pY88 system shows significantly deviant RMSD values over time compared to Control, strongly suggesting Y88 phosphorylation most perturbs WPD loop dynamics, closely followed by pY-all. Earlier experimental studies on ligand-free Yersinia PTP and the enzyme complexed with oxyanions^74,75^ have suggested the significance of a potential hydrogen bond formation between W354 and R409 residues (numbering in Yersinia PTP) in facilitating the closure of the WPD-loop. Simulation studies by Kamerlin et al.^76,77^ on the WPD loop dynamics in PTP1B have also implicated the distance between residues W179 and R221 (numbering in PTP1B) as a good measure to characterize open/closed state of WPD loop. Along similar lines, we have monitored the distance between W197 and R237 (numbering in PTP-PEST system and equivalence obtained from sequence and structure alignment of PTP1B and PTP-PEST systems) to probe the role of tyrosine phosphorylation on the WPD loop dynamics and the results are shown in Figure 7. The results are in line with the RMSD values of WPD loop with tyrosine phosphorylation and show that in pY88 system, the significant increase in distance between W197 and R237 further supports the effect of pY88 on WPD loop dynamics. To further understand WPD loop conformational sampling and phosphorylation effects, Principle Component Analysis (PCA) of an expanded 193-209 WPD loop set was performed. The spread of points in a PC1 vs PC2 plot defines explored conformational space. The largest spreads for the WPD loop occur for pY88 and pY-all systems (Figure S6), agreeing with the RMSD results. pY64 and pY64+pY88 show similar scatter plots (Figure S6(b,d)). The results also agree with lost D199-K142 and D199-237 interactions (Figure 6(b-c)), indicating Y88 phosphorylation affects WPD loop conformation and flexibility.

**FIG. 7.**
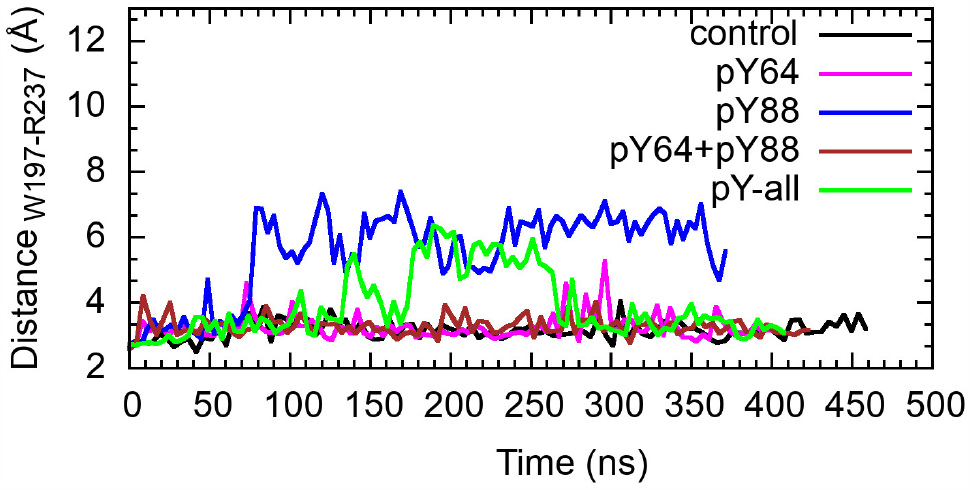
Time evolution of distance between W197(O) and R237(N) in (a) control (b) pY64 (c) pY88 (d) pY64+pY88 (e) pY-all PTP-PEST systems simulation length scale.

Next we investigated the impact of tyrosine phosphorylation on the catalytic site size, focusing on distances between crucial residues lining the site for substrate binding and interaction. These include D199, K142, Q278, and p/Y64 (depending on Y64 phosphorylation), with measurements of distances K140 - Q278 and Y64 - D199 depicted in Figure 8(a). The corresponding distance histogram is presented in Figure 8(b). A noteworthy shift in these distances is evident for the pY88 system, strongly indicating a potential opening of the binding pocket. An increase in distances, though smaller compared to pY88, is also observed for the pY-all system. To delve deeper, we revisited the network analysis, specifically examining the optimal paths between residue Y88 and the entire WPD loop (197-199) in both the control and the pY88 system. The results are displayed in Figure S7. The analysis strongly suggests that phosphorylation of the Y88 residue alters the optimal path, connecting Y88 and the entire WPD loop in a single path, in contrast to the control system. In the pY88 system, the pY88 residue exhibits strong interactions not only with the WPD loop but also with flanking residues, both sequentially and spatially. This observation implies a robust allosteric link between the Y88 residue and the WPD loop. Collectively, these analyses demonstrate that tyrosine phosphorylation significantly influences access and the structure of the catalytic site, particularly in the case of pY88, potentially impacting substrate binding to PTP-PEST.

**FIG. 8.**
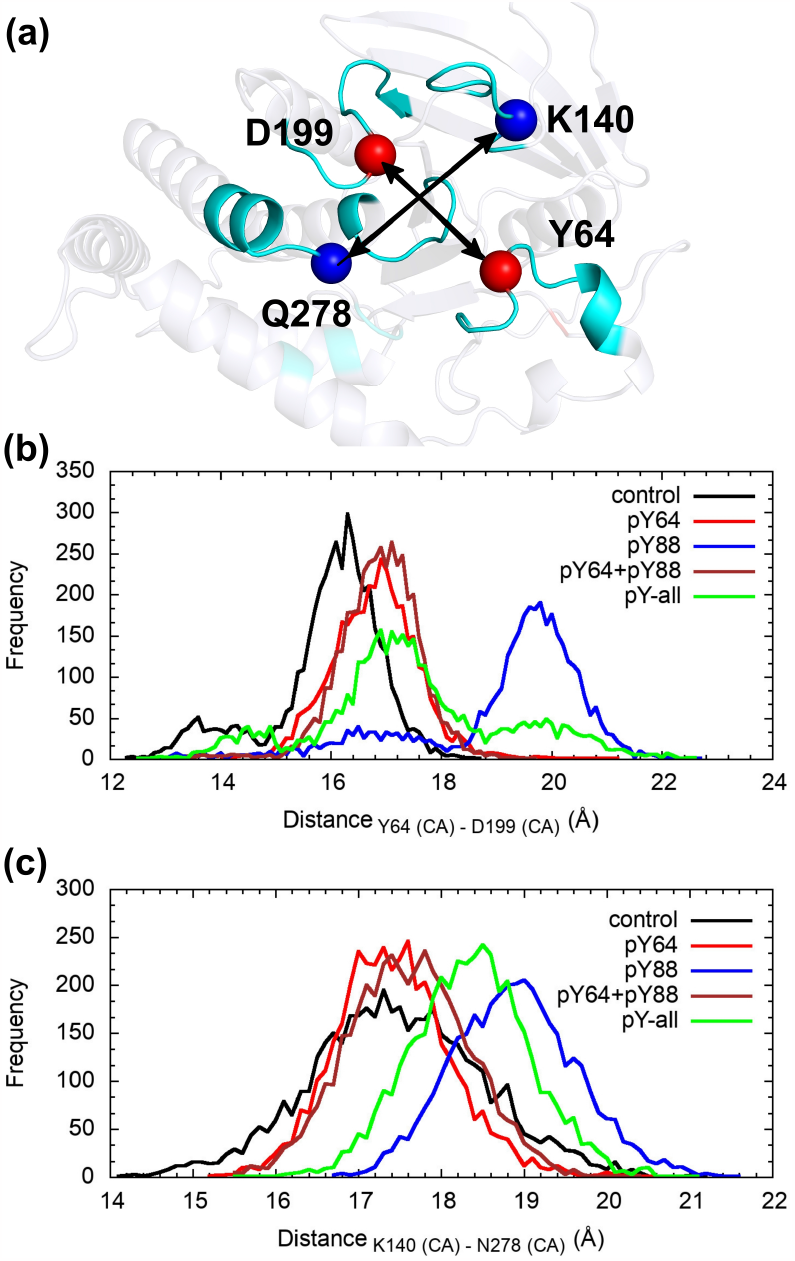
(a) cartoon representation of binding groove where the C*α* atoms of relevant residues D199, Y64, Q278 and K140 are shown as spheres. Frequency distribution of (a) K140 - N278 and (c) Y64 - D199 of control, pY64, pY88, pY64+pY88 and pY-all PTP-PEST systems

### E. Tyrosine phosphorylation on substrate binding

Previous analyses have shown phosphorylation(s) can affect substrate binding to the phosphorylated PTP-PEST proteins. Small catalytic site environment changes can impact substrate binding^3,78^. We studied binding of two substrates, HER2 hepta-peptide and Src-derived mutant peptide inhibitor, to various equilibrated Control and phosphorylated systems. The substrate structures were taken from a PTPN18 catalytic domain-HER2 complex (PDB: 4GFU) and a PTP1B-Src inhibitor complex (PDB: 3ZMQ). Notably, the HER2 peptide is longer than the Src one, allowing us to probe effects of catalytic site depth and structure changes on binding. Active site docking of the crystal structure ligands to control and phosphorylated complexes was performed in Autodock Vina^69^ and binding affinities calculated in PRODIGY^71,72^ (Table III; IC details in Tables S1 and S2). The HER2 and Src binding conformations varied across all phosphorylated PTP-PEST proteins compared to control (Figure 9(a)).

**TABLE 3.** Binding affinity of SRC derived peptide and HER2 derived peptide to control and phosphorylated PTP-PEST systems calculated using PRODIGY webserver.

| Protein-peptide complex | $\Delta G$ (kcal/mol) | Protein-peptide complex | $\Delta G$ (kcal/mol) |
| --- | --- | --- | --- |
| control-HER2-complex | -7.4 | control-SRC-complex | -7.0 |
| pY64-HER2-complex | -6.9 | pY64-SRC-complex | -6.5 |
| pY88-HER2-complex | -8.8 | pY88-SRC-complex | -6.9 |
| pY64+pY88-HER2-complex | -7.9 | pY64+pY88-SRC-complex | -7.1 |
| pY-all-HER2-complex | -8.0 | pY-all-SRC-complex | -6.2 |

**FIG. 9.**
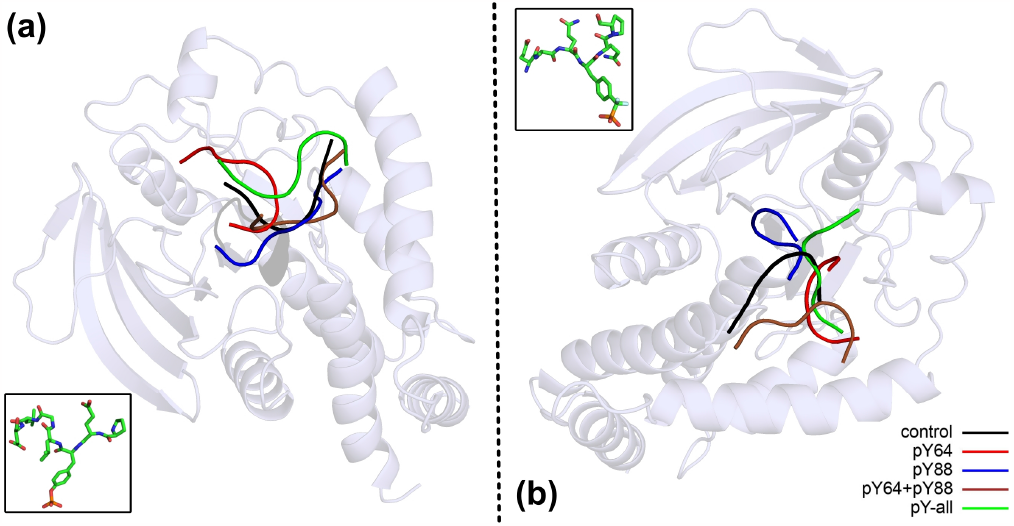
Cartoon representation showing the binding conformation of (a)HER2-derived peptide and (b)SRC-derived peptide along with the stick representation of respective bound peptide.

The change in peptide binding affinities differs for the two peptides across phosphorylated systems. Binding energy calculations show the HER2-derived peptide has significantly greater affinity for pY88 compared to pY64+pY88 and pY-all systems versus Control (Table III), while pY64 shows a moderate decrease (Table III). In pY88, the bound HER2 peptide adopts a more extended, buried conformation within the wider catalytic pocket, likely due to WPD loop flexibility changes seen previously. For pY64, Y64 phosphorylation seems to negatively affect HER2 binding, with N-terminal residues moving away, no longer interacting, potentially reducing affinity. Consequently, in double phosphorylated pY64+pY88, the modifications additively allow a more extended peptide conformation and increased affinity compared to Control, but less than pY88 alone. Interestingly, in pY-all, the four phosphorylations have an overall positive effect enabling extended peptide binding with higher affinity than Control.

The shorter SRC peptide (6 aa) seems to bind very differently than the longer HER2 peptide (9 aa), possibly due to the importance of length in determining binding conformation and protein-peptide interactions. The SRC peptide can access more conformations with similar affinity in the binding groove due to its shorter length. This is supported by the very different bound conformations across systems (Figure 9(b)). For SRC, there is no significant affinity change for the phosphorylated PTP-PEST systems (Figure 9(b)). The binding conformation is such that although the phosphate-containing residue faces into the pocket, the remaining residues point away, likely reducing pY64 interactions versus Control. In SRC bound to pY-all, the peptide flips orientation compared to Control (Figure 9(b)). While new contacts form with pY-all, this drastic conformational difference may reduce binding affinity. These docking studies show PTP-PEST phosphorylation(s) have a significant, differential effect on substrate binding that depends on substrate size. This may be substrate-specific and relate to the portion interacting with phosphorylated PTP-PEST.

## IV. DISCUSSION AND CONCLUSIONS

The regulation of protein activity is fundamental for controlling cell behavior and maintaining normal function. Protein phosphorylation, a post-translational modification, plays a pivotal regulatory role. In particular, the reversible phosphorylation of proteins by kinases and phosphatases can significantly influence various cellular processes. Although relatively rare, tyrosine phosphorylation holds special significance in eukaryotic signaling, alongside the more common serine and threonine phosphorylation. Phosphorylation is known to modulate protein stability, enzymatic activity, and substrate affinity^79–82^. Previous studies identified serine phosphorylation sites on PTP-PEST, but here we focused on understanding the potential for tyrosine phosphorylation regulation.

Our study protein, PTP-PEST, is an 88 kDa cytosolic protein belonging to the protein tyrosine phosphatase (PTP) family. PTP-PEST regulates various physiological processes including cell adhesion, migration, immune response, and apoptosis. Despite its importance, the potential for tyrosine phosphorylation and implications for PTP-PEST function were largely unexplored. We began by confirming PTP-PEST can indeed undergo tyrosine phosphorylation. Using bioinformatics prediction tools, we identified specific PTP-PEST tyrosine residues as putative phosphorylation sites. Notably, Y64 and Y88 were identified as highly conserved, phosphorylation-prone residues. Mutational studies indicated Y64 significance for PTP-PEST catalytic activity. Y64 is situated in the pTyr-recognition loop, playing a role in binding pocket depth. It has been reported as essential for PTP-PEST activity^33^. Y88 lies within an important hydrophobic cluster in the structure. Y194 is positioned in the catalytic WPD loop and Y283 in the Q-loop, close to glutamines crucial for the catalytic reaction. This intriguing discovery suggested PTP-PEST’s catalytic domain could be dynamically regulated by tyrosine phosphorylation.

To gain insights into the global effects of phosphorylation, we conducted comprehensive molecular dynamics simulations. Our results revealed changes in protein flexibility, inter-residue interactions, and conformational dynamics, particularly in PTP-PEST’s key catalytic loops. These findings indicated tyrosine phosphorylation can impact structural dynamics, potentially influencing substrate recognition and binding. Further investigations examined the role of phosphorylation in loop dynamics, focusing on the flexible WPD-loop. Our analysis showed phosphorylation, especially at Y88, influenced WPD-loop flexibility. As this loop acts as a gate to the active site, this dynamic alteration may have important implications for substrate accessibility to the catalytic pocket. Additionally, we explored phosphorylation effects on catalytic pocket size, showing it could lead to binding pocket dimensional variations. These changes might affect substrate interactions with PTP-PEST, potentially influencing binding affinity and specificity. To understand the functional consequences of these structural insights, we conducted docking studies using PTP-PEST homolog structures with bound substrates. These revealed tyrosine phosphorylation could have substrate-specific effects on binding affinities and bound peptide conformational changes. Importantly, phosphorylation of residues like Y88 seemed to enhance binding of certain substrates, underlining the regulatory potential of tyrosine phosphorylation in PTP-PEST function.

In summary, our comprehensive study including experimental, bioinformatics and extensive atomistic simulations, sheds light on the previously unexplored realm of tyrosine phosphorylation in PTP-PEST. We have demonstrated that this reversible modification can profoundly affect structural dynamics and functional aspects of the protein. These findings highlight the intricate regulatory mechanisms governing cellular processes involving PTP-PEST. We have shown tyrosine phosphorylation profoundly impacts PTP-PEST, influencing structure, dynamics, and substrate binding. Understanding PTP-PEST’s regulatory mechanisms through tyrosine phosphorylation opens new avenues for exploring its role in cellular processes and potential therapeutic interventions.

## V. ACKNOWLEDGEMENTS

All the simulations in this work have been carried out on clusters Annapurna and Nandadevi at The Institute of Mathematical Sciences, Chennai, India. Experimental work was funded through the ‘Centre of Excellence (CoE) in Molecular Medicine’ grant awarded to Madhu-lika Dixit.

## Notes

### Competing Interest Statement

The authors have declared no competing interest.

