## Supplementary Data for "Role of Tyrosine Phosphorylation in PTP-PEST"

| Y-position | Score |
|---|---|
| 42 | 0.439 |
| 48 | 0.508 |
| 64 | 0.949 |
| 88 | 0.984 |
| 98 | 0.418 |
| 103 | 0.451 |
| 124 | 0.432 |
| 146 | 0.635 |
| 150 | 0.627 |
| 173 | 0.441 |
| 190 | 0.371 |
| 194 | 0.916 |
| 219 | 0.385 |
| 246 | 0.672 |
| 283 | 0.92 |
| 301 | 0.475 |
| 387 | 0.951 |
| 409 | 0.416 |
| 560 | 0.516 |
| 733 | 0.386 |

**Table S1.** NetPhos Analysis of tyrosine phosphorylation sites across PTP-PEST. Scores above a threshold of 0.7 are highlighted in red.

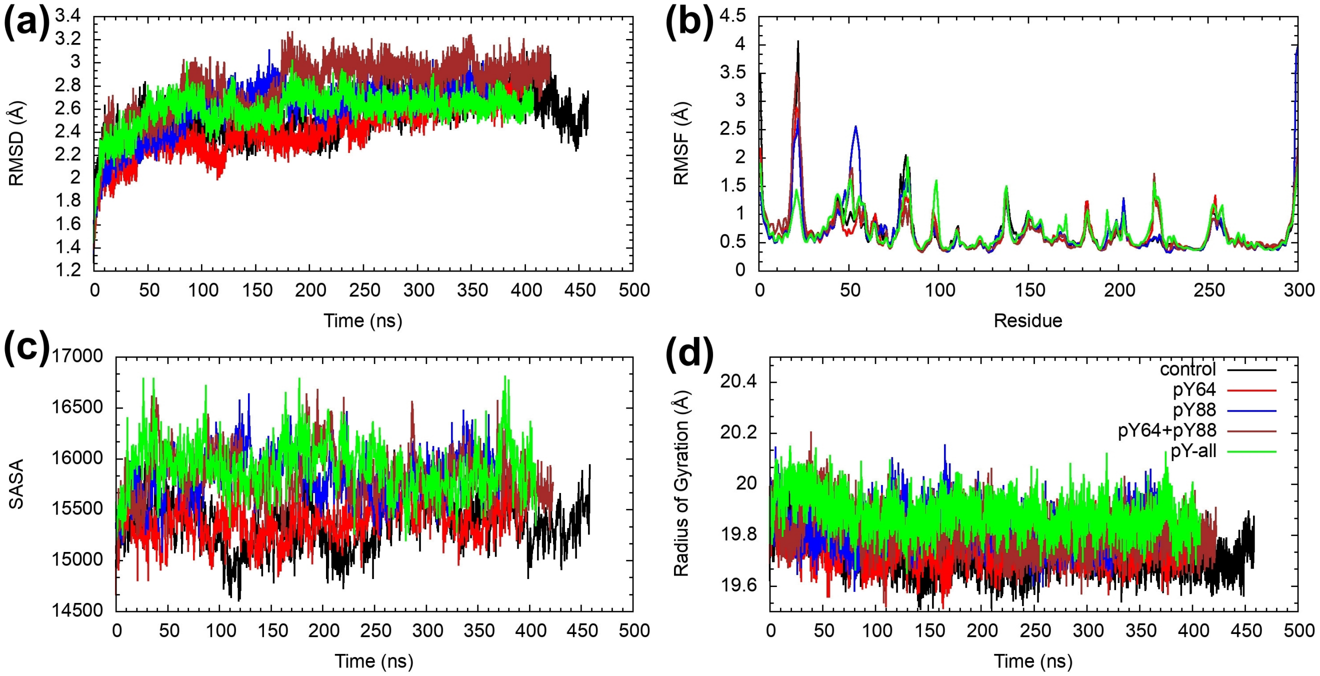

**Figure S1.** Time evolution of (a) Root Mean Square Deviation (RMSD) of PTP-PEST , (b) Root Mean Square Fluctuation (RMSF) of PTP-PEST averaged over the last 25ns , (c) Radius of Gyration (Rg) of PTP-PEST and (d) Solvent Accessible Surface Area (SASA) of PTP-PEST

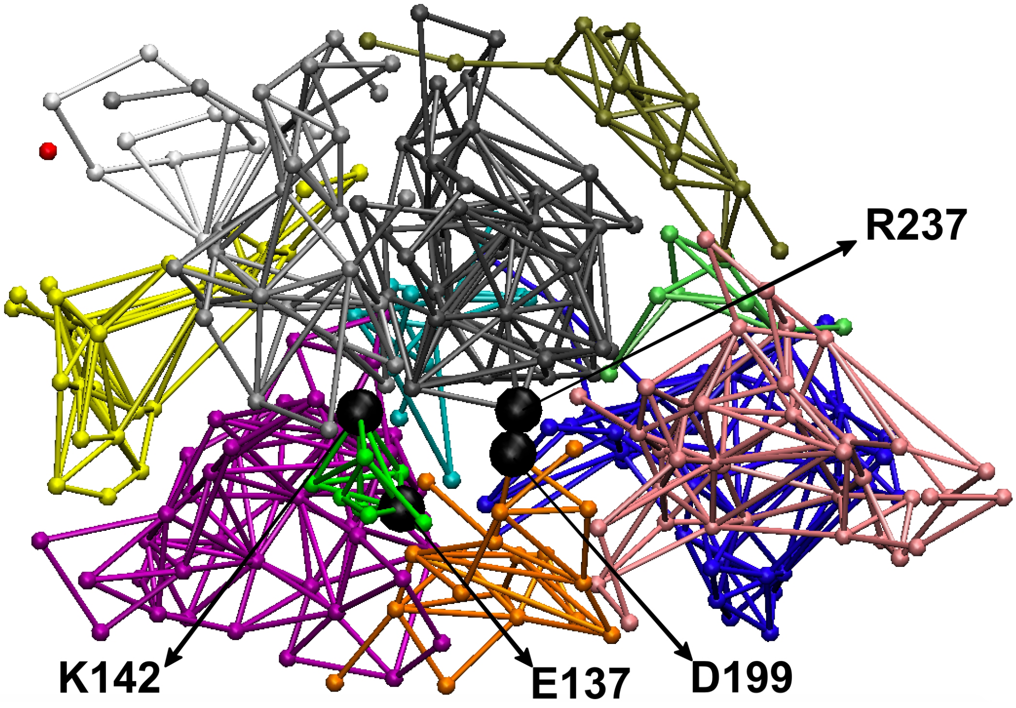

**Figure S2.** Network analysis over the last 50ns of pY-all-PTP-PEST system

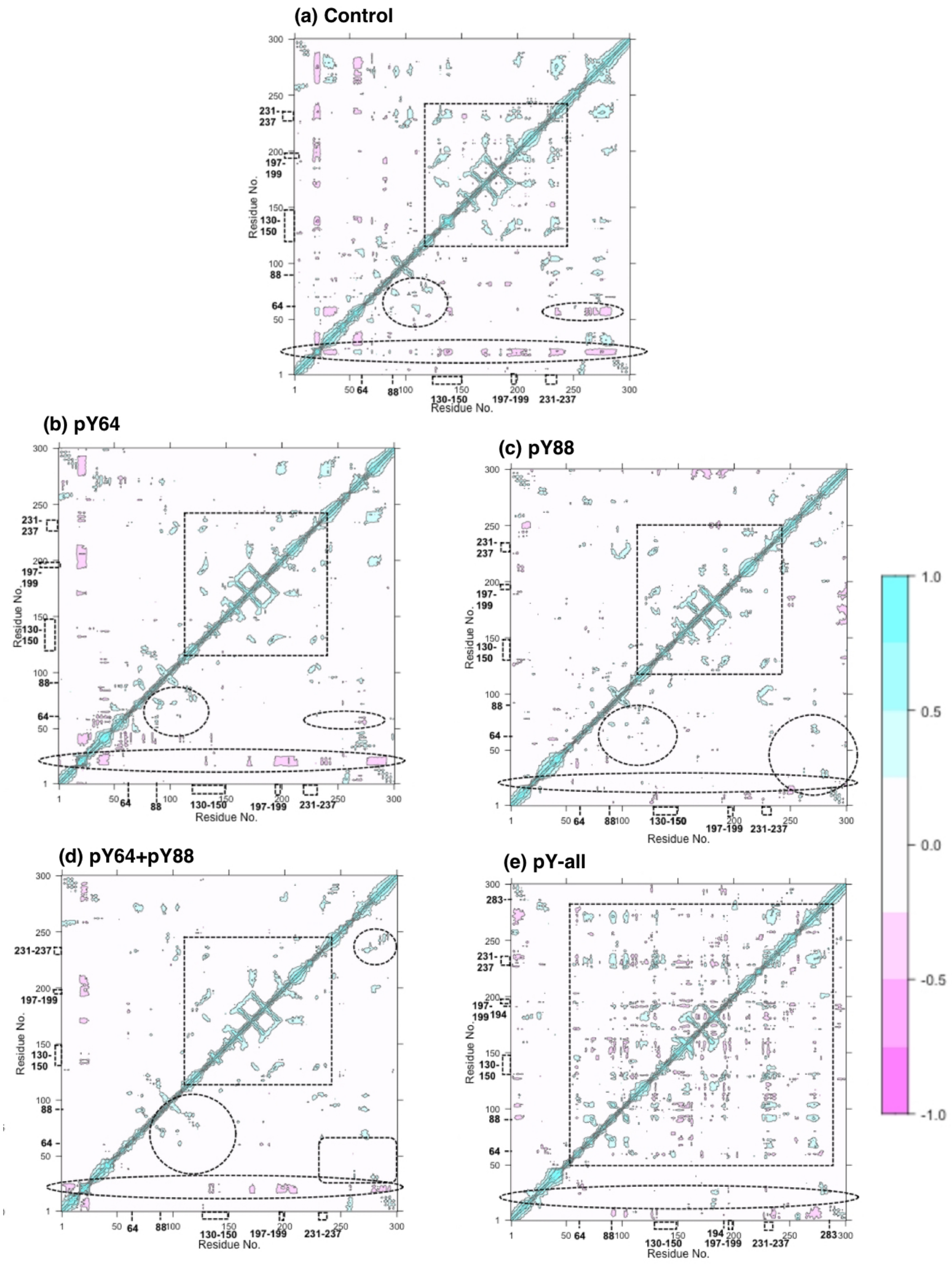

**Figure S3.** Cross correlation plots calculated over last 50 ns of simulation for (a) control, (b) pY64, (c) pY88, (d) pY64+pY88 and (e) pY-all PTP-PEST systems. The color bar indicates positive(cyan) to negative(pink) correlations. Residues of interest are specifically numbered and the regions of correlation which differ from the control system, as different tyrosines are phosphorylated are also indicated as dotted surfaces.

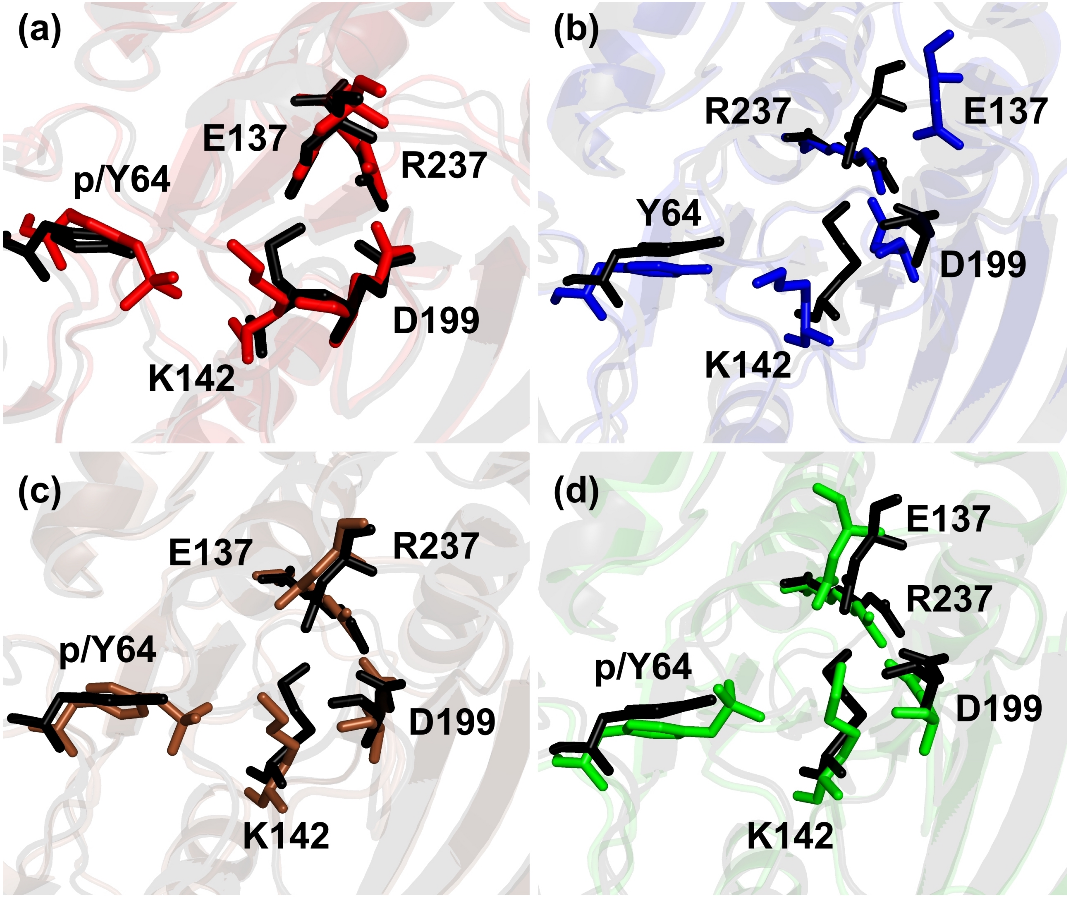

**Figure S4.** Graphical representation of residues E137, R237, Y64 , D199 and K142 in control (black), (a) pY64(red), (b) pY88(blue), (c)pY64+pY88(brown) and (d)pY-all(green) systems. The phosphorylated and unphosphorylated Y64 is marked as p/Y64 respectively

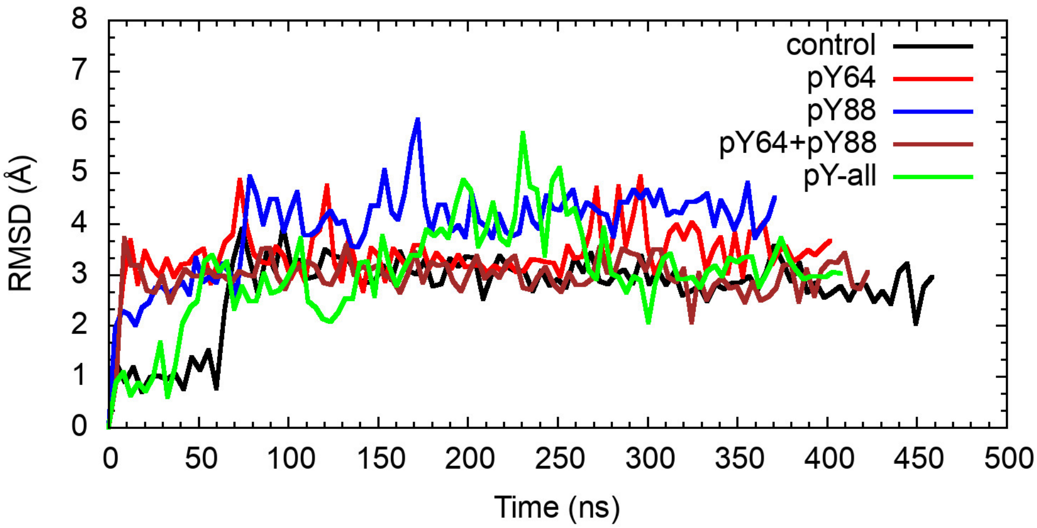

**Figure S5.** Time evolution of RMSD of C$\alpha$ atom of residues 197-203 (WPD-loop) for control, pY64, pY88, pY64+pY88 and pY-all PTP-PEST systems.

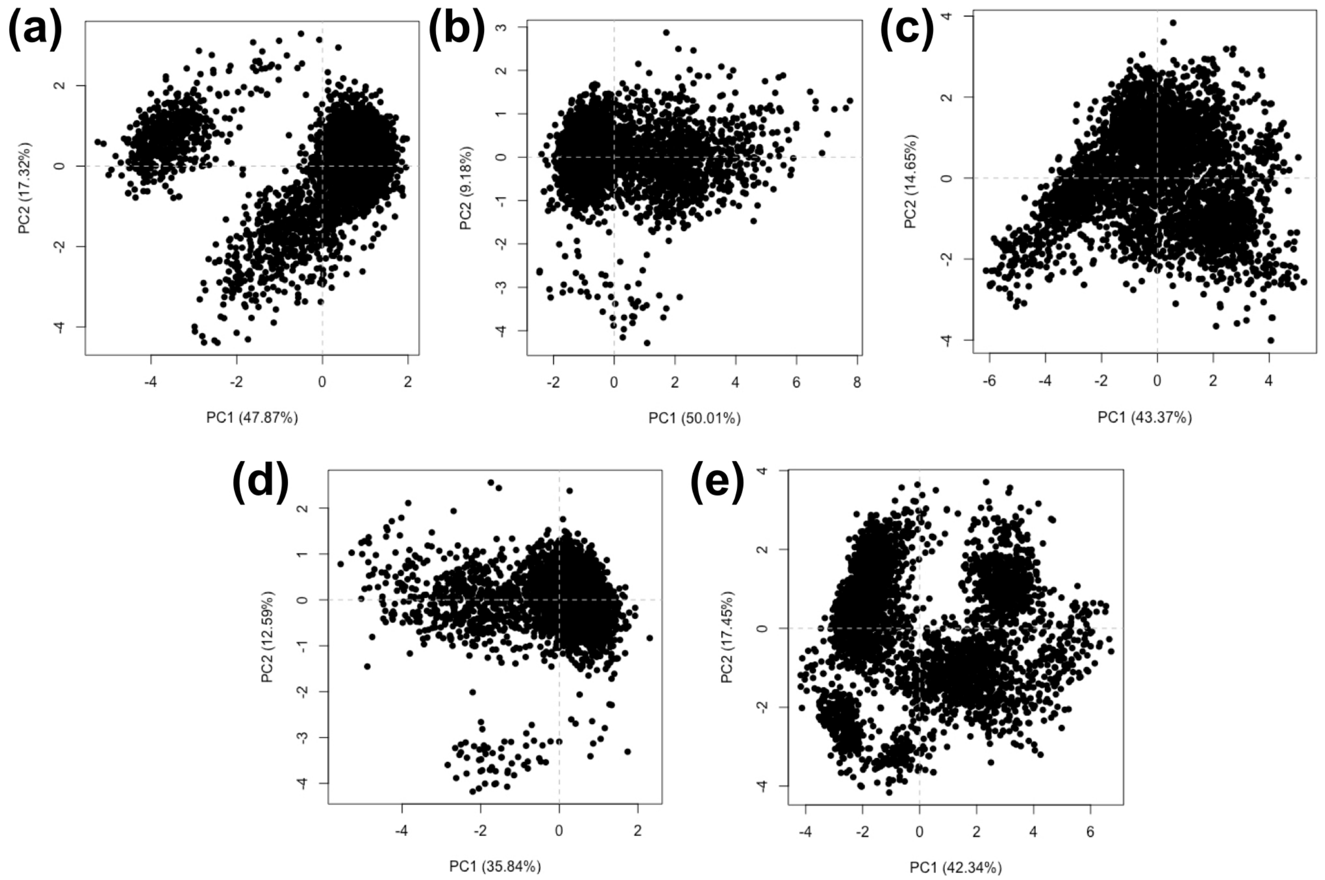

**Figure S6.** PC1 v/s PC2 plots from principal component analysis of WPD loop (193-209) of (a) control (b) pY64 (c) pY88 (d) pY64+pY88 (e) pY-all PTP-PEST systems over the stretch of simulation.

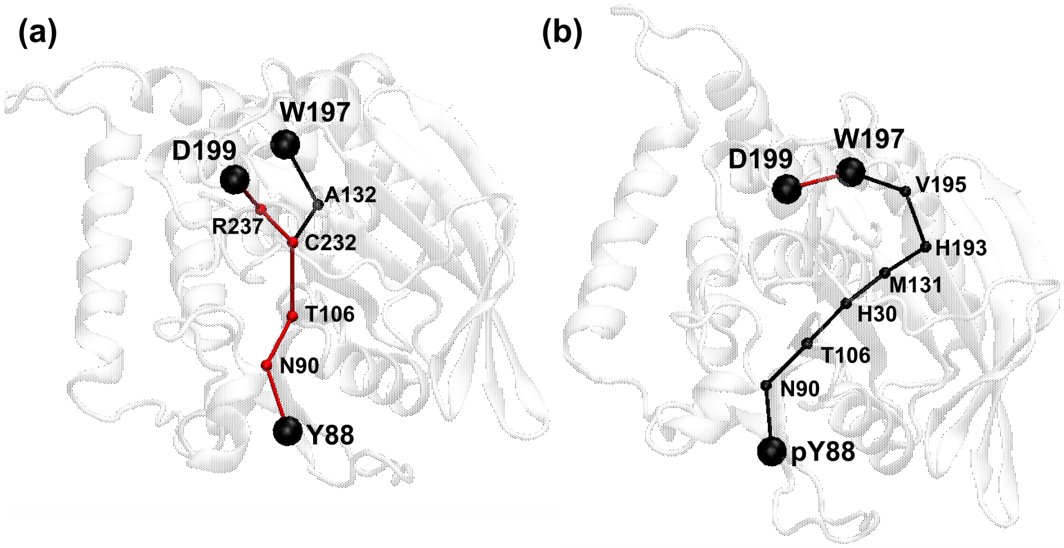

**Figure S7.** Optimal path calculated between Y88 and W197(black) and Y88 and D199 (red) in (a) control and (b) pY88 PTP-PEST systems using NetworkView plugin of VMD.

| Position | Code | Kinase | PSP | Score | Cutoff |
|---|---|---|---|---|---|
| **48** | Y | TK | TPSQDSDYINANFIK | 0.2958 | 0.2874 |
| 48 | Y | TK/ALK | AVQTKEQYELVHRAI | 0.3408 | 0.2737 |
| 48 | Y | TK/ALK | HPAGGIHYEMCIECP | 0.2919 | 0.2737 |
| **64** | Y | TK/Abl | KYRTEKIYPTATGEK | 0.2484 | 0.2458 |
| **88** | Y | TK/Abl | GVYGPKAYVATQGPL | 0.2906 | 0.2458 |
| 88 | Y | TK/Abl | HPAGGIHYEMCIECP | 0.2497 | 0.2458 |
| 88 | Y | TK/Axl | TPSQDSDYINANFIK | 0.3001 | 0.2459 |
| 88 | Y | TK/Axl | MISLMRKYQEHEDVP | 0.2669 | 0.2459 |
| 88 | Y | TK/Csk | GVYGPKAYVATQGPL | 0.3208 | 0.258 |
| 88 | Y | TK/EGFR | TPSQDSDYINANFIK | 0.3289 | 0.2774 |
| **103** | Y | TK/EGFR | VWQDNDRYHPKPVLH | 0.3327 | 0.2774 |
| 103 | Y | TK/EGFR | HPAGGIHYEMCIECP | 0.32 | 0.2774 |
| 103 | Y | TK/Eph | ENVKKNRYKDILPFD | 0.3309 | 0.2719 |
| **146** | Y | TK/Eph | HPAGGIHYEMCIECP | 0.3323 | 0.2719 |
| **219** | Y | TK/FGFR | KYRTEKIYPTATGEK | 0.2884 | 0.2788 |
| 219 | Y | TK/FGFR | GRKKCERYWPLYGED | 0.2843 | 0.2788 |
| 219 | Y | TK/FGFR | HPAGGIHYEMCIECP | 0.3278 | 0.2788 |
| 219 | Y | TK/FAK | TPSQDSDYINANFIK | 0.3053 | 0.2718 |
| **283** | Y | TK/Fer | KYRTEKIYPTATGEK | 0.2781 | 0.2464 |
| **387** | Y | TK/Fer | GVYGPKAYVATQGPL | 0.2821 | 0.2464 |
| 387 | Y | TK/PDGFR | TPSQDSDYINANFIK | 0.3502 | 0.3187 |
| **733** | Y | TK/PDGFR | MISLMRKYQEHEDVP | 0.4709 | 0.3187 |
| 733 | Y | TK/Ret | MISLMRKYQEHEDVP | 0.3031 | 0.252 |
| 733 | Y | TK/Ret | HPAGGIHYEMCIECP | 0.3078 | 0.252 |
| 733 | Y | TK/Src | MISLMRKYQEHEDVP | 0.3122 | 0.2703 |
| 733 | Y | TK/Trk | HPAGGIHYEMCIECP | 0.318 | 0.2925 |
| 733 | Y | TK/VEGFR | TPSQDSDYINANFIK | 0.515 | 0.3642 |
| 733 | Y | TK/VEGFR | VWQDNDRYHPKPVLH | 0.3729 | 0.3642 |

**Table S2.** GPS 6.0 Analysis of tyrosine phosphorylation sites across PTP-PEST. Predicted tyrosine phosphorylation sites were obtained after the highest threshold was applied. The scores obtained for each kinase that possibly phosphorylates these tyrosines are given along with their respective cutoffs.

| Protein-peptide complex | ΔG (kcal/mol) | Kd (M) | ICs charged-charged | ICs charged-polar | ICs charged-apolar | ICs polar-polar | ICs polar-apolar | ICs apolar-apolar |
|---|---|---|---|---|---|---|---|---|
| control-HER2-Complex | -7.4 | 3.8E-06 | 4 | 3 | 14 | 0 | 4 | 4 |
| pY64-HER2-Complex | -6.9 | 8.4E-06 | 10 | 3 | 10 | 0 | 2 | 2 |
| pY88-HER2-Complex | -8.8 | 3.5E-07 | 11 | 5 | 16 | 0 | 7 | 6 |
| pY64+ pY88 -HER2-Complex | -7.9 | 1.7E-06 | 6 | 6 | 14 | 0 | 5 | 3 |
| pY-all-HER2-Complex | -8.0 | 1.4E-06 | 4 | 6 | 16 | 0 | 6 | 2 |

**Table S3.** Binding affinity , dissociation constant (Kd) and number of intermolecular contacts (ICs) at the interface of HER2 derived peptide and control and phosphorylated PTP-PEST systems calculated using PRODIGY webserver.

| Protein-peptide complex | ΔG (kcal/mol) | Kd (M) | ICs charged-charged | ICs charged-polar | ICs charged-apolar | ICs polar-polar | ICs polar-apolar | ICs apolar-apolar |
|---|---|---|---|---|---|---|---|---|
| control-SRC-complex | -7.0 | 7.7E-06 | 4 | 6 | 8 | 0 | 4 | 4 |
| pY64-SRC-complex | -6.5 | 1.6E-05 | 3 | 7 | 7 | 3 | 6 | 3 |
| pY88-SRC-complex | -6.9 | 8.2E-06 | 3 | 9 | 6 | 4 | 9 | 4 |
| pY64+pY88-SRC-complex | -7.1 | 5.9E-06 | 5 | 6 | 7 | 4 | 8 | 2 |
| pY-all-SRC-complex | -6.2 | 3E-05 | 4 | 5 | 7 | 2 | 3 | 1 |

**Table S4.** Binding affinity , dissociation constant (Kd) and number of intermolecular contacts (ICs) at the interface of SRC derived peptide and control and phosphorylated PTP-PEST systems calculated using PRODIGY webserver.
